# Ceylon cinnamon and its major compound Cinnamaldehyde can limit overshooting inflammatory signaling and angiogenesis *in vitro*: implications for COVID-19 treatment

**DOI:** 10.1101/2021.06.16.448642

**Authors:** Kurt Lucas, Maximilian Ackermann, Anna Lena Leifke, William W. Li, Ulrich Pöschl, Janine Fröhlich-Nowoisky

**Affiliations:** Max Planck Institute for Chemistry, Multiphase Chemistry Department, Mainz, Germany; Institute of Pathology and Molecular Pathology, Helios University Clinic Wuppertal, University of Witten/Herdecke, Wuppertal, Germany; Institute of Functional and Clinical Anatomy, University Medical Center of the Johannes Gutenberg-University Mainz, Mainz, Germany; The Angiogenesis Foundation, Cambridge, Massachusetts, United States of America

**Keywords:** Ceylon cinnamon extract (CCE), Cinnamaldehyde (CA), Coronavirus disease 2019 (COVID-19), Dexamethasone, Toll-like receptor 4 (TLR4)

## Abstract

Overshooting immune reactions can occur during inflammatory responses that accompany severe infections, such as COVID-19. Cytokines, damage-associated molecular patterns (DAMPs), and reactive oxygen and nitrogen species can generate positive feedback loops of inflammation, leading to long-term complications such as vascular endothelialitis, thrombosis, endothelial dysfunction, neurological impairments, and chronic fatigue. Dexamethasone can limit inflammation by inhibiting the activation of pro-inflammatory transcription factors. High dose dexamethasone, however, has undesirable side effects. Here, we show that Ceylon cinnamon and its major compound cinnamaldehyde can mitigate inflammatory signaling *in vitro*. Cinnamaldehyde interferes with the dimerization of toll-like receptor 4 (TLR4), which can be activated by DAMPs like HSP60 and HMGB1. Our results suggest that supplementary treatment with Ceylon cinnamon may allow administration of lower doses of dexamethasone to avoid high dose steroid side effects. Moreover, preliminary results indicate that Ceylon cinnamon modulates angiogenesis, which is a reactive phenomenon in COVID-19.

## Introduction

Coronavirus disease 2019 (COVID-19) causes an acute inflammatory response in organs with frequent long-term effects. Ackermann et al. 2020 described massive pulmonary vascular endothelialitis, thrombosis, and angiogenesis in COVID-19 patients (1). Long-term complications include neurological disturbances, kidney and myocardial disorders, as well as chronic fatigue (2–5). COVID-19 infection is also characterized by excessive immune reactions, which are frequently referred to as the ‘cytokine storm’ (6, 7). Various cytokines are involved in this overshooting immune reaction, including TNF, IFN-γ, IL-1β, IL-2, IL-4, IL-6, IL-8, and CRP (8–10). The transcription factor nuclear factor kappa-light-chain-enhancer of activated B cells (NF-κB, p50/p65) is also heavily involved in inflammatory processes (Fig. 1), regulating the expression of hundreds of inflammatory genes, including the pro-inflammatory cytokines IL-1β and TNF (11, 12), generating a positive feedback loop (Fig. 1). Moreover, damage-associated molecular patterns (DAMPs) like heat shock protein 60 (HSP60), high mobility group box 1 (HMGB1), and calprotectin (S100A8/A9 hetero tetramer) can be released during immune reactions elicited by COVID-19 infection (13–16). These DAMPs can activate pattern recognition receptors such as TLR2, TLR4, and RAGE (14, 16–19) and generate additional positive feedback via NF-κB, as has been shown for TLR4 (Fig. 1), causing further cytokine release. Additionally, high amounts of reactive oxygen and nitrogen species (ROS/RNS) are produced during inflammation (20). As we previously reported, ROS/RNS can substantially potentiate the TLR4 stimulation potential of DAMPs *in vitro*, suggesting that these species can probably further escalate inflammatory processes (19, 21) (Fig.1).

**Figure 1.**
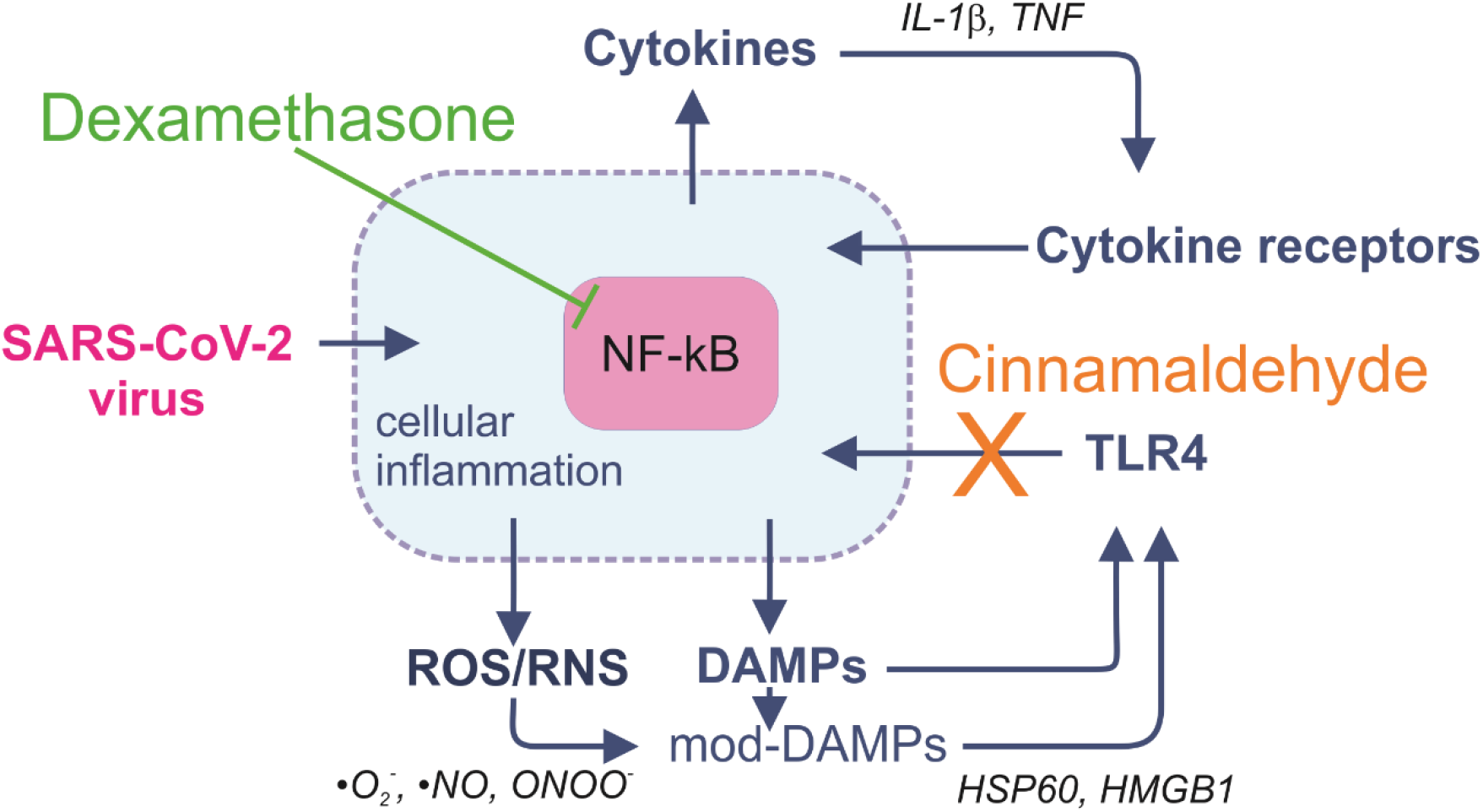
Positive feedback loops amplifying the magnitude of inflammatory signaling. Pro-inflammatory cytokines cause activation of NF-κB, which amplifies their production, resulting in a positive cytokine feedback loop. In addition, DAMPs can be released and activate NF-κB via TLR4 and other pattern recognition receptors, generating a further feedback loop. Moreover, interactions with ROS/RNS can form chemically modified DAMPs (mod-DAMPs) with enhanced TLR4-stimulating potential, additionally fueling inflammation. Dexamethasone inhibits activation of NF-κB, whereas cinnamaldehyde, the major active compound of ethanolic Ceylon cinnamon extract, inhibits activation of TLR4, thus disrupting the feedback loops.

One of the clinical goals for the treatment of COVID-19 is to mitigate an excessive inflammatory response to SARS-CoV-2 infection. Dexamethasone, a corticosteroid which acts on NF-κB (22, 23) is an effective therapeutic intervention for moderate to severe disease (4, 24), but it can cause unwanted side effects in high doses. TLR4 could be another potential target for treatment of COVID-19 since it plays an important role in the DAMP-driven positive feedback loops (Fig. 1). Cinnamaldehyde (CA), the major active compound of Ceylon cinnamon (*Cinnamomum verum*) extract (CCE) (25–27) can inhibit the activation of TLR4 by preventing receptor dimerization (28). As shown in an animal model, the concurrent intake of Cinnamon can also mitigate the side effects of dexamethasone (29). We hypothesized that Ceylon cinnamon or cinnamaldehyde might be well suited as a supplementary treatment for COVID-19 to lower the risk of the ‘cytokine storm’ and allow administration of reduced dosages of dexamethasone to mitigate steroid-induced side-effects (30).

In this study, we compared CCE and CA with dexamethasone in the suppression of TLR4-dependent cytokine mRNA production and IL-8 protein release *in vitro*. Further, we present preliminary data showing that CCE inhibits tube formation, which serves as a cellular model for angiogenesis.

We do not recommend the use of Cassia cinnamon or its extracts, due to its possibly high content of liver-toxic coumarin (31).

## Materials and Methods

### Ceylon cinnamon extract and cinnamaldehyde

Ceylon cinnamon extract (CCE) was prepared by extraction from Ceylon cinnamon powder (*Cinnamomi ceylanici cortex*, Caesar & Loretz GmbH, Hilden, Germany) with 70% ethanol (Sigma-Aldrich Chemie GmbH, Taufkirchen, Germany) in a 1:5 ratio and incubation for 10 days at room temperature, protected from light and shaken once daily. After the incubation period, the CCE was filtered through a 0.22-μM pore size PES membrane filter unit (Corning Inc., Corning, USA) under sterile conditions and stored in brown glass bottles at room temperature protected from light. In a previous study, a CCE concentration of 0.3% was determined to be optimal for the treatment of THP-1 cells (26), with higher concentrations resulting in cytotoxic effects; lower concentrations were, on the other hand, less potent in preventing inflammation (26). CCE was diluted 1:10 in Dulbecco’s Phosphate Buffered Saline (DPBS -/-) (Thermo Fisher Scientific Inc., Waltham, USA) and cells were treated with final concentrations of 0.2% and 0.3% CCE.

A cinnamaldehyde (CA, Sigma-Aldrich) stock solution of 20 mg/mL was freshly prepared in absolute ethanol (Sigma-Aldrich). For cell culture experiments, we diluted the CA stock solution in DPBS -/- to concentrations of 1 mM and 10 mM.

### Dexamethasone

A dexamethasone (Sigma-Aldrich) stock solution (1 mg/mL in 70% ethanol) was freshly prepared for each experiment. THP-1 cells were treated with final dexamethasone concentrations of 3.2 μM and 6.4 μM according to Menacher et al., 2017 (32).

### Cultivation and treatment of THP-1 monocytes

Cell culture experiments were performed with human THP-1 acute monocytic leukemia cells (ATCC, LGC Standards GmbH, Wesel, Germany). Cells were grown in Roswell Park Memorial Institute (RPMI) 1640 medium (Thermo Fisher Scientific) supplemented with 10% heat-inactivated fetal bovine serum (FBS superior; Biochrom, Berlin, Germany), 0.05 mM 2-mercaptoethanol (Sigma-Aldrich) and 100 U/mL penicillin-streptomycin (Thermo Fisher Scientific) at 37 °C in a 5% CO_2_ humidified atmosphere.

The THP-1 cells were seeded in 96-well V-bottom cell culture plates (Greiner Bio-One GmbH, Frickenhausen, Germany) at a density of 4×10^4^ cells in 100 μL growth medium per well and allowed to settle for 1 h. In duplicate, cinnamon extract or vehicle control (70% ethanol) was added to the cells in final concentrations of 0.2% and 0.3%, and cells were pre-incubated for 2 h. Cinnamaldehyde was added in concentrations ranging from 10 μM up to 70 μM, also including a vehicle control (100% ethanol), followed by pre-incubation in the same way. After pre-incubation, TLR4 activation was stimulated by adding lipopolysaccharide (LPS-EB from *E. coli* 0111:B4; InvivoGen, Toulouse, France) in a final concentration of 50 ng/mL. Cells were incubated with LPS for 4 h. Supernatants were used to determine IL-8 release using an enzyme-linked immunosorbent assay (ELISA; BD, Heidelberg, Germany). Cell viability was determined following overnight incubation with alamarBlue™ cell viability reagent (Thermo Fisher Scientific), which was added to the cells in a concentration of 10%. After incubation, the fluorescence intensity was measured with a Synergy™ NEO HTS multi-mode microplate reader (Biotek Instruments GmbH, Bad Friedrichshall, Germany) using excitation and measurement wavelengths of 560 nm and 590 nm, respectively. Two independent experiments were performed.

For qPCR analysis, cells were seeded in 6-well cell culture plates at a density of 4×10^5^ cells/mL and treated in triplicate with 0.3% cinnamon extract, 3.2 μM or 6.2 μM dexamethasone, and CA in a range of 20 μM–50 μM plus vehicle control as described above. LPS incubation was carried out for 1 h and 4 h. After incubation, cells were separated from supernatants by centrifugation (500 × *g*, 5 min), lysed in RLT buffer (RNeasy Mini Kit;Qiagen, Hilden, Germany) and stored at −80 °C for RNA extraction and real-time quantitative PCR. Two independent experiments were performed on different days.

### IL-8 ELISA

Interleukin-8 secretion of treated THP-1 monocytes was quantified by the ELISA OptEIA™ Set for human interleukin-8 (BD) following the manufacturer’s protocol with optimized wash buffer composition (50x dilution). The cell supernatants were diluted in a range of 1:2–1:20 and tested in triplicate. The absorbance wavelength of 450 nm and reference wavelength of 570 nm were determined on a Synergy NEO plate reader. Using Gen5 software (Biotek), the reference wavelength values were subtracted from the absorbance wavelength values. Based on measured standard values, a standard curve was constructed. IL-8 cytokine production was calculated according to the standard curve and respective dilution factors. Statistical analysis and corresponding graphs were generated with Prism 9 software (GraphPad Software, San Diego, USA).

### RNA extraction and qPCR

Real-time quantitative PCR (qPCR) was performed to quantify mRNA expression of IL1-β, IL-8, and TNF. Extraction of total RNA from cells was performed using the RNeasy Mini Kit (Qiagen) according to the manufacturer’s protocol. Purified RNA was eluted with 30 μL RNase-free water. Total RNA yield was determined by measuring absorbance at 260 nm, 280 nm, and 320 nm using Take3 Trio Micro-Volume Plates (Biotek) in combination with the Synergy NEO plate reader. All samples were adjusted to an RNA concentration of 50 ng/μL. Single-strand cDNA was synthesized from 500 ng RNA per sample using the High-Capacity cDNA Reverse Transcription Kit (Thermo Fisher Scientific) following the standard protocol. cDNA (2.1 ng) served as template for a qPCR reaction using SsoAdvanced™ Universal SYBR^®^ Green Supermix (Bio-Rad Laboratories, Hercules, USA) and forward and reverse primers in a final concentration of 333 nM each. Primer sequences were designed with Primer-BLAST software (NCBI) (Table 1). Peptidylprolyl Isomerase A (PPIA) and TATA-Box Binding Protein (TBP) served as reference genes (33). The following thermal cycling protocol was used: 98 °C for 30 s followed by 40 cycles of 98 °C for 10 s, and 60 °C for 25 s. Data analysis was performed with CFX Manager software 3.1 (Bio-Rad) using the 2-^ΔΔCT^ method to calculate gene expression. For qPCR, all samples were tested in technical duplicates and arithmetic mean and standard deviation values were calculated. Prism 9 software (GraphPad Software) was used for statistical analysis.

**Table 1.**
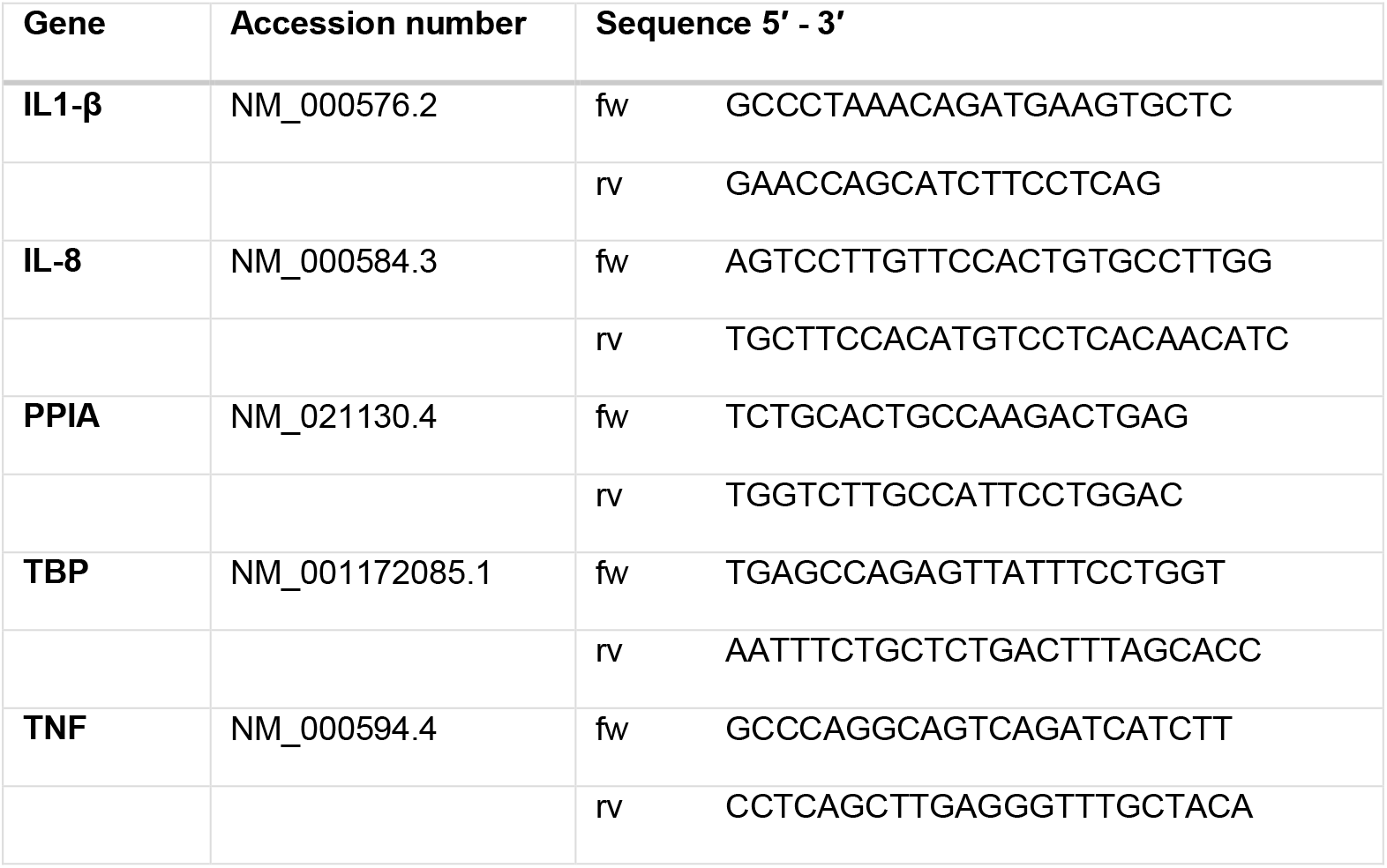
PCR primer sequences. Primer pairs used for qPCR; cytokines: interleukin 1 beta (IL-1β), interleukin 8 (IL-8), tumor necrosis factor (TNF); reference genes: peptidylprolyl isomerase A (PPIA) and TATA-box binding protein (TBP).

### Cultivation of HUVEC Cells

HUVEC cells (Lonza) were cultivated in endothelial cell medium (EGM-2; Lonza) containing 2% fetal bovine serum (FBS) and vascular endothelial growth factor (VEGF) for rapid proliferation. To analyze tube formation, the cells were first grown (seeding density 2 × 10^4^cells/mL) in 6-well plates and then transferred to collagen-coated bottom plates without basement membrane extract (CELLCOAT-coated; Biocompare, South San Francisco, CA, USA). The cells were stained with Calcein-AM (#17783, Sigma, Taufkirchen, Germany) and inspected with a fluorescence microscope (Olympus, Hamburg, Germany) using excitation and emission wavelengths of 496 nm and 520 nm.

### Endothelial cell tube formation assay

A commercial assay system was used (Thermo Fisher Scientific, Waltham, MA, USA) to determine endothelial cell tube formation; the assay was performed as described in the instructions from the manufacturer and following a published procedure(34). In this system, HUVEC cells (from Lonza, Basel, Switzerland) were cultivated in EGM-Plus Growth Medium (with 5 mM glucose), containing supplements (35) at 37 °C with 5% CO_2_. For the experiments, cells at passage <11 were used. The matrix, formed from collagen/basement membrane extract (Geltrex; Thermo Fisher Scientific), was layered into 12 well-plates (Corning/Costar-Sigma, Taufkirchen, Germany). The dishes were overlaid with 1 × 10^5^ cells/well in 400 μL of conditioned medium. Tube formation was checked during the first 10 h using a reflection electron microscope (REM). Electron microscopy was performed with a scanning electron microscope (ESEM) using an ESEM XL-30 apparatus (Philips, Eindhoven; Netherlands).

## Results and Discussion

To evaluate toxicity, an alamarBlue™ assay was performed to determine cell viability (Fig. 2). Ceylon cinnamon extracts at concentrations of 0.2% and 0.3% elicited no toxic effects in THP-1 cells, which is in accordance with previous results (26). At concentrations of up to 40 μM, cinnamaldehyde showed no toxic effects in THP-1 monocytes, whereas concentrations of CA of 50 μM and higher resulted in reduced cell viability. Dexamethasone concentrations of 3.2 μM and 6.4 μM elicited no cytotoxic effects.

**Figure 2.**
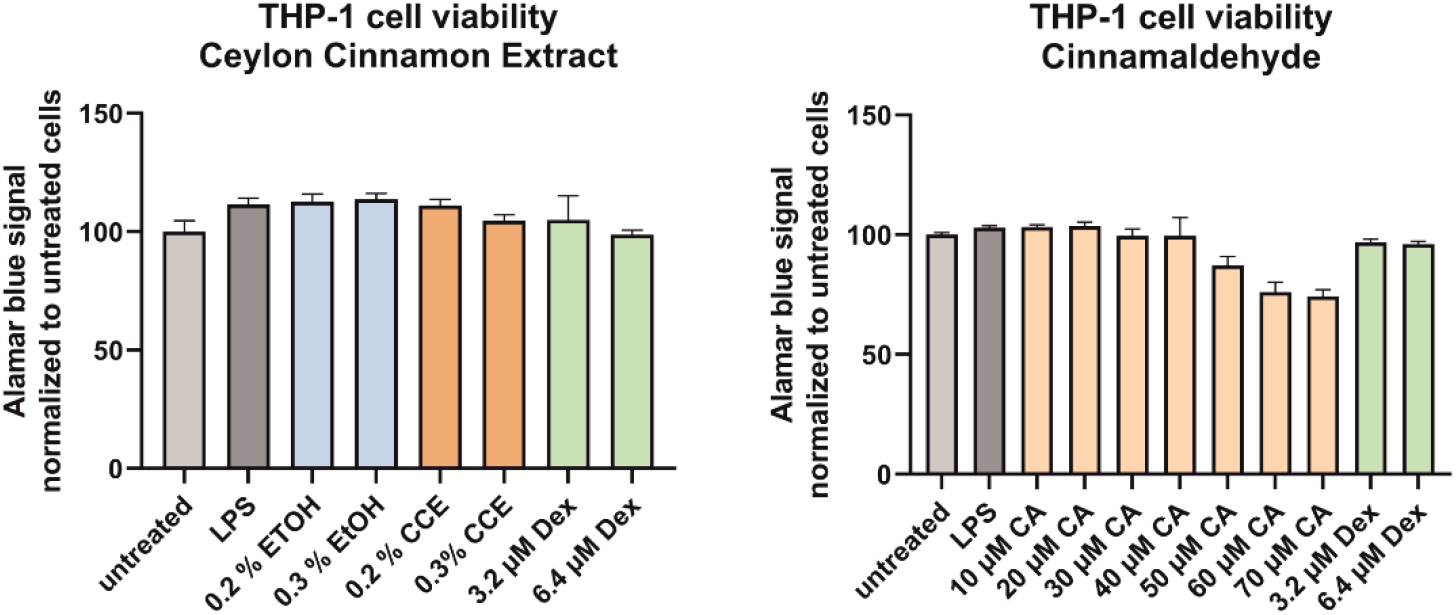
Cell viability. THP-1 monocytes were incubated with Ceylon cinnamon extract (CCE) at final concentrations of 0.2% and 0.3%, equivalent ethanol concentrations, dexamethasone (Dex), or with various concentrations of cinnamaldehyde (CA). After 2 h, cells were challenged or not with LPS in a final concentration of 50 ng/mL and incubated for 4 h. Thereafter, the alamarBlue™ assay was performed to determine cell viability. Arithmetic mean values and standard deviations based on four measurement for each column; both experiments were repeated with very similar results on different days.

### Expression of cytokine genes

We next tested the efficacy of CCE and CA in inhibiting IL-1β, IL-8, and TNF mRNA expression and IL-8 protein release in THP-1 monocytes (Fig. 3, Fig. 4). In previous studies, we determined that a concentration of 0.3% CCE is most effective in THP-1 cells (26, 27). To confirm dose dependency, we also included a concentration of 0.2% CCE in the mRNA expression experiments. For CA, we tested concentrations of 20 μM, 30 μM, 40 μM, and 50 μM CA. We compared CCE and CA with two concentrations of dexamethasone (3.2 μM and 6.4 μM) as a reference standard anti-inflammatory drug.

**Fig. 3.**
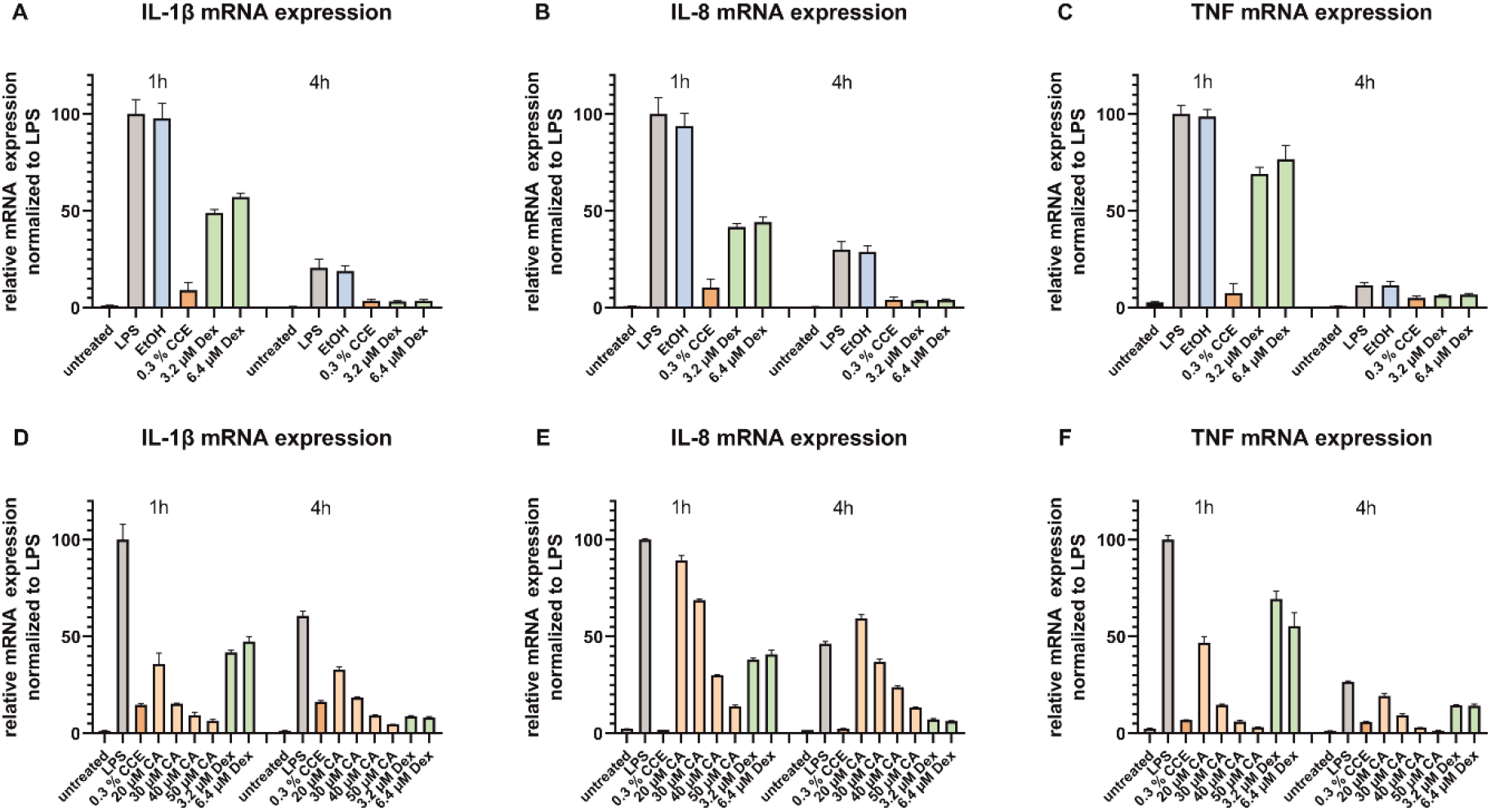
Effects of Ceylon cinnamon extract (CCE) and cinnamaldehyde (CA) on cytokine mRNA expression. Relative IL-1β, IL-8, and TNF mRNA abundance after different treatments, normalized to LPS stimulation. Cells were pre-incubated with 0.3% CCE, different concentrations of CA or dexamethasone (3.2 μM and 6.4 μM) for 2 h. Then, LPS (final conc. 50 ng/mL) was added and cells were incubated for a further 1 h or 4 h. mRNA was isolated and quantified by qPCR. Arithmetic mean values and standard deviations of two independent experiments performed on different days as triplicates, measured in technical duplicate.

**Fig 4.**
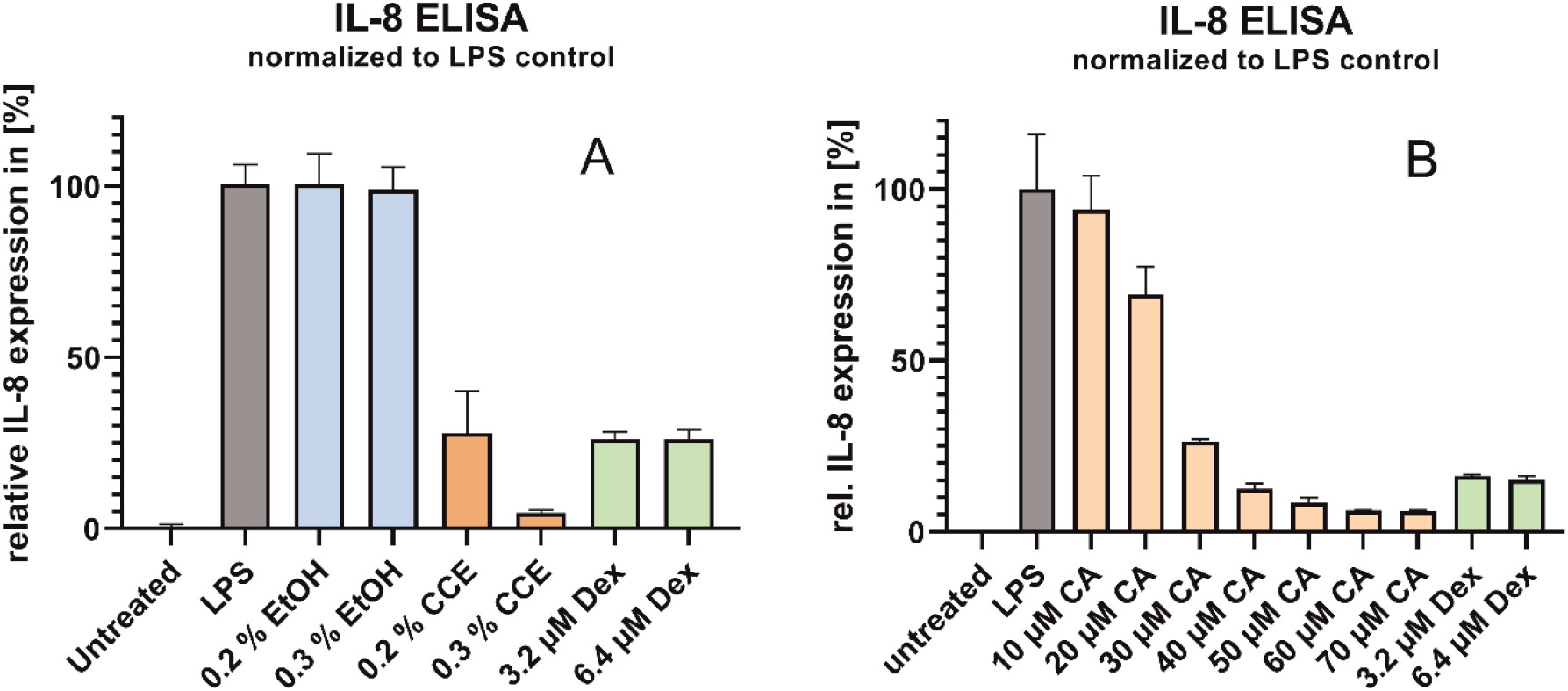
IL-8 protein expression. Relative protein expression of IL-8 in THP-1 cells challenged for 4 h with 50 ng/mL LPS and treated with CCE, equivalent concentration of ethanol (A), CA, or dexamethasone (B). Cells were pre-incubated with the respective treatment for 2 h prior to LPS challenge. Arithmetic mean values and standard deviations of two independent experiments performed on different days as biological duplicates, measured in technical duplicate.

### Inhibition of cytokine mRNA expression by CCE

The mRNA expression of all tested NF-κB-regulated cytokines was strongly enhanced 1 h after stimulation with 50 ng/mL LPS with sharp decreases in the induction by 4 h (Fig. 3). The used ethanol concentrations had no impact on gene expression. Treatment with 0.3% CCE drastically inhibited the LPS-stimulated increases in the mRNA expression of all three cytokines. In contrast, the inhibitory effects of 3.2 μM and 6.4 μM dexamethasone after 1 h were relatively lower, but similar to CCE after 4 h. This suggests that the effects of dexamethasone on inflammatory signaling pathways are not immediate (36).

Cinnamaldehyde showed a clear dose dependence in reducing the mRNA expression of all three cytokines examined (Fig. 3). This was expected, since the cinnamaldehyde in CCE interferes with TLR4 receptor dimerization and thus inhibits the activation of NF-κB by LPS (28). This can explain the faster action of cinnamaldehyde compared to dexamethasone, because inflammatory TLR4 signaling is blocked already outside the cell. As shown in Figure 2, a CA concentration of 50 μM elicited cytotoxic effects, which additionally influences gene expression. Concentrations of 30 μM CA and 3.2 μM dexamethasone were similarly potent in suppressing inflammatory signaling. In these experiments, the effect of 30 μM CA on IL-1β and TNF expression was comparable to that of CCE, but the effect was less pronounced for IL-8. Overall, whole CCE seems to be more efficient than cinnamaldehyde in the suppression of mRNA expression of inflammatory genes. We also observed this phenomenon in a previous study; one possible explanation might lie in the synergistic effects of CA with *p*-cymene, cinnamyl alcohol, and cinnamic acid, which are additional active compounds of CCE (27).

For quantification of the effects of CCE, CA, and dexamethasone on IL-8 cytokine release, THP-1 monocytes were pre-incubated with 0.2% or 0.3% CCE, equivalent concentrations of ethanol, or with 3.2 μM or 6.4 μM dexamethasone (Fig. 4A), and seven different concentrations of CA (Fig. 4B). LPS was added in a final concentration of 50 ng/mL, and the cells were incubated for a further 4 h. Supernatants were collected and analyzed by IL-8 ELISA. The baseline IL-8 expression of untreated cells was set to zero, while the IL-8 expression of the LPS-treated cells was set to 100% in both experimental settings. The addition of 0.2% and 0.3% ethanol had no effect on IL-8 protein expression. CCE in a concentration of 0.2% had a similar effect to 3.2 μM or 6.4 μM dexamethasone (Fig 4A). CCE at a concentration of 0.3% was more effective for the suppression of IL-8 than either of the dexamethasone concentrations used. CA at concentrations of 30 μM and 40 μM inhibited TLR4-dependent IL-8 expression to an extent comparable to the dexamethasone concentrations used (Fig. 4B).

Taken together, CCE and CA can dampen the activation of TLR4-dependent pathways during inflammation. This suggests that cinnamon compounds may also reduce the ROS/RNS-stimulated, DAMP-mediated activation of TLR4 during severe inflammation (Fig. 1).

### In vitro angiogenesis assay

Reactive angiogenesis is another complication present in SARS-CoV-2-infected organs in COVID-19. The tube formation assay in HUVEC cells is a common cellular model for angiogenesis (37). We carried out preliminary tube formation assay experiments *in vitro* to determine whether CCE can also mitigate this complication of COVID-19.

If HUVEC cells are cultivated onto solubilized and subsequently solidified basement membrane extract matrix, they start to form a tube-like network (Fig. 5A). This morphogenetic pattern is caused by a sprouting of endothelial cells, which is routinely used as a surrogate for angiogenesis. The temporal analysis of this morphogenetic process by fluorescence light microscopy revealed an inhibitory effect of 0.1% CCE after 3.5 h and 6 h hours (Fig. 5B), whereas higher dosages (0.5% and 1.0% cinnamon, Fig. 5C and 5D) showed direct cytotoxic effects. The analysis of this sprouting process by scanning electron microscopy (SEM) showed an altered cell morphology in response to 0.1% CCE (Fig. 5F). Higher dosages resulted in cellular damage and apoptosis (Fig. 5G and 5H).

**Fig 5.**
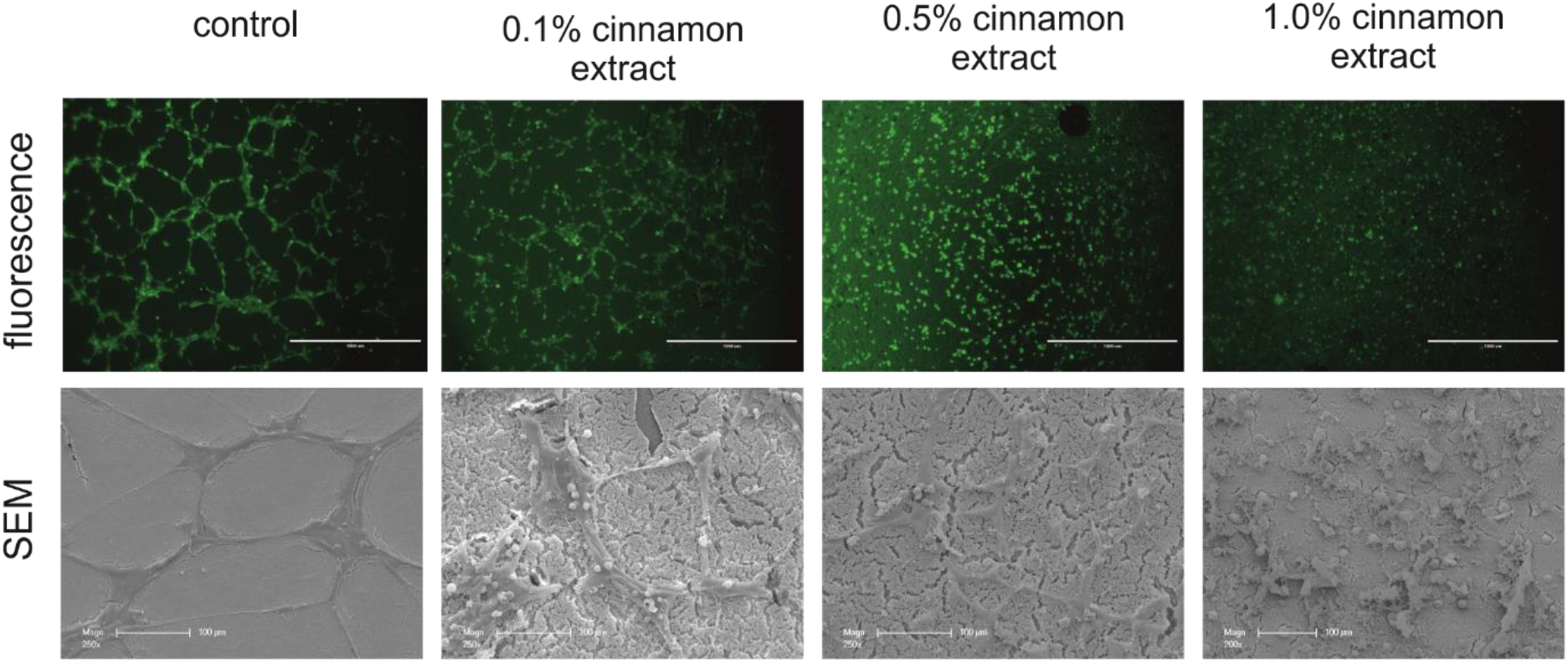
*In vitro* formation of tubes. HUVEC cells were grown on solubilized basement membrane extract; fluorescence light microscopy (Fig. 5A–5D). After seeding 1 × 10^5^ cells per well in 400 μL of medium, tube formation was recorded over the course of 6 h. Where indicated, the cells were inspected at time 0 (seeding), after 3.5 h, and 6 h. Prior to microscopic inspection, the cells were stained with calcein-AM and inspected under fluorescence light; A: control; B: 0.1% Ceylon cinnamon extract (CCE); C: 0.5% CCE; D: 1.0% CCE. Fig. 5E–5H: HUVEC cell formation at higher magnification after a total incubation period of 16 h; analysis by scanning electron microscopy (SEM). In contrast to controls, 0.1% CCE resulted in an attenuation of tube formation, whereas 0.5% and 1.0% CCE elicited cytotoxic effects in HUVEC cells.

The inhibition of angiogenesis by CCE and CA is well documented in the literature (38–43). Suprisigly one studie find a angigenesis promoting effect for CA in the context of wouldhealing (44).

### Side effects of corticosteroids

Despite the clinical efficacy of dexamethasone for COVID-19 patients, unwanted side-effects are known for corticosteroids, e.g. steroid psychosis (45), hyperglycemia, especially in diabetic patients (46), and secondary bacterial pneumonia or invasive fungal infection from immunosuppression (47). Hypothetically, cinnamon or cinnamaldehyde should act synergistically with dexamethasone. A combination of both may thus allow the administration of a reduced dosage of dexamethasone. As shown in an animal model, the concurrent intake of cinnamon can mitigate the side-effects of dexamethasone (48). Cinnamon can also positively influence insulin resistance, lipid metabolism, and glucose transport (48, 49).

### Dosage form and dosage

Ethanolic cinnamon extract is much more suitable than cinnamon powder for cell culture experiments, since, in contrast to the powder form, it can be applied quantitatively by pipetting directly into the culture medium. In contrast, the majority of clinical studies on cinnamon have used encapsulated cinnamon bark powder (30). Since almost no reports describing ethanolic CCE for human use have been published to date, future studies on COVID-19 patients should use the encapsulated powder. Typically, doses of encapsulated cinnamon vary between 500 mg and 3 g per day for adults (50–54). However, to the best of our knowledge, no study administering cinnamon alone or in combination with dexamethasone for COVID-19 has been published to date. Therefore, clinical studies are needed before the treatment of COVID-19 patients with cinnamon or its compounds can be recommended. The Ceylon cinnamon should be of pharmaceutical grade as described here.

Since dexamethasone and cinnamon act to dampen inflammation via distinct pathways, we suggest that these compounds are likely to have synergistic effects when administered concomitantly.

## Conclusions

Cinnamon has been used as a medicine for thousands of years in traditional medical practices (55). Its anti-inflammatory effects are well documented. The inhibition of TLR4 dimerization, in particular, is an important anti-inflammatory mechanism (28). During the so called ‘cytokine storm’, not only cytokines but also ROS/RNS and DAMPs contribute to amplification of inflammation (56). Our results indicate that dexamethasone, which is now being used to dampen excessive inflammation in COVID-19, could be combined with a Ceylon cinnamon preparation. In contrast to dexamethasone, which inhibits the activity of the pro-inflammatory transcription factor NF-κB (22), cinnamon can suppress the secondary activation of TLR4 by DAMPs (28). These differences in modes of action suggest the possibility of synergistic effects. Further studies may pave the way towards clinical studies of cinnamon derivatives for the treatment of COVID-19.

Our results suggest that intake of Ceylon cinnamon in combination with dexamethasone may prevent the ‘cytokine storm’ and help to reduce the dosage of dexamethasone and mitigate its side effects. Preliminary results indicate that cinnamon may also reduce angiogenesis. Note, however, that Ceylon cinnamon should not be substituted by Cassia cinnamon, which may comprise high amounts of liver-toxic coumarin (31).

## Abbreviation list

CCE: Ceylon cinnamon extract
CE: cinnamaldehyde
CRP: C-reactive protein
DAMP: damage-associated molecular pattern
Dex: dexamethasone
HMGB1: high mobility group box 1 protein
HSP60: heat shock protein 60
HUVEC: human umbilical vein endothelial cells
IFN-γ: interferon gamma
IL-1β: interleukin 1 beta
RAGE: receptor for advanced glycation endproducts
ROS/RNS: reactive oxygen and nitrogen species
TLR4: toll-like receptor 4
TNF: tumor necrosis factor

## Author contributions

KL, AM, ALL and UP designed and conducted research; KL, AM, ALL and JF-N analyzed data; KL, AM, ALL, JF-N, UP and WL wrote and edited the manuscript; KL, AM, JF-N and UP had primary responsibility for final content.

